# A rational approach for multicopy delta integration in *Saccharomyces cerevisiae* targeting a conserved Ty1 LTR sequence

**DOI:** 10.1101/2025.09.26.678811

**Authors:** Gustavo Seguchi, Beatriz de Oliveira Vargas, Gonçalo Amarante Guimarães Pereira, Fellipe da Silveira Bezerra de Mello

## Abstract

Genomic integration is a foundation in yeast-based cell factories, ensuring phenotypic stability and adequate gene dosage to sustain desired metabolic fluxes. Retrotransposon-associated sequences are attractive genomic integration targets for metabolic engineering, as they are present in multiple copies throughout the genome and their disruption should not jeopardize the cell. In *S. cerevisiae*, multiple strategies have been proposed to target Ty elements, but little attention has been given to the homology sequence itself. Through analysis of all long terminal repeats in *S. cerevisiae*, we propose a novel consensus δ sequence for efficient multicopy gene integration. Reducing homology length to favor a more conserved sequence greatly increases the number of potential genomic integration sites – reaching 85 targets in this work. The designed integration sequence was validated by introducing a non-native xylose fermentation pathway into this yeast, achieving a single-step simultaneous integration of 15 xylose isomerase copies, enabling efficient ethanol production in the engineered strain.

## Introduction

The yeast *Saccharomyces cerevisiae* is a widely used microbial platform in biotechnology (1). Its uses range from the production of sophisticated biopharmaceuticals such as insulin and vaccine antigens (2) to the production of commodities like ethanol on massive scales (3). The versatility of this microorganism stems from its amenability to genetic engineering. Complex heterologous pathways comprising dozens of steps can be assembled in this cell factory, steering native metabolism towards desired products (4,5). Although an extensive toolbox for epissomal expression in yeast is available to create tailored recombinant cells (6–8), antibiotics or nutritional supplementation are required for plasmid maintenance (9). Such an approach, however, becomes impractical at industrial scales. On the other hand, chromosomal integration offers a viable strategy for stable gene expression, as it preserves the desired modifications without the need for selective pressure while providing consistent expression levels by introducing a defined number of gene copies.

The challenge then becomes how and where to insert the genes of interest (GOIs). The yeast once again proves itself useful. Whereas mammalian cells require elaborate techniques to allow insertion of foreign DNA (10), yeast naturally favor the homologous recombination (HR) pathway for double strand break (DSB) DNA repair (11). This allows straightforward incorporation of DNA into yeast genome by flanking them with homologous sequences to the desired chromosomal site (12). There are several suitable sites where GOIs can be inserted. Common examples are the genes involved in nutrient metabolism (*URA3, LEU2, MET15, HIS3* etc.), the homing endonuclease (HO) *locus* as well as rationally selected “safe harbors” (13–17). However, when a pathway involves multiple genes or when multiple copies of a gene are needed to achieve the desired expression level, each successive round of integration in these sites requires the design and cloning of tailored homology arms. In this sense, retrotransposons represent well suited sites for successive genetic manipulation.

These genomic parasites spread through the genome by transcribing themselves into RNA molecules that are then reverse-transcribed and inserted into distinct locations (18). In *S. cerevisiae*, the prototypical retrotransposons are the Ty (Transposon yeast) elements, namely the families Ty1-Ty5. The most prominent of them are the Ty1 elements, structured as two open reading frames flanked by two 334 bp long terminal repeats (LTRs), also called delta sequences (19). The *Saccharomyces* genome database (SGD) contains 31 full Ty1 copies and 299 solo delta sequences (20), formed by recombination between the LTRs as a mechanism to prevent further transpositions (21,22).

Many methods have been devised to target GOI multicopy integration into these sites (23–33). All these approaches rely on an arbitrarily selected delta sequence as a homology template. However, this may not represent the optimal strategy, as a single delta sequence might not adequately reflect the genomic diversity. Since most delta sequences are scars of ancient transposition events, these sequences are subject to accumulating mutations over evolutionary time. They can undergo deletions, leading to significantly reduced sizes (<100 bp), insertions, leading to significantly increased sizes (>500) or minor mutations that do not significantly alter their size (22). Therefore, a highly diverged delta sequence would be a poor homology template, leading to low integration efficiency.

To provide a rationally designed delta sequence for multicopy integration in *S. cerevisiae*, this study aims to select, clone and validate a consensus delta sequence representing the most conserved region among all annotated sequences in the yeast genome. A bioinformatic pipeline was developed to determine the best candidate target, which was amplified from genomic DNA and cloned into a plasmid allowing straightforward multicopy integration of any GOI. The strategy was validated by integrating an *Orpinomyces* sp. xylose isomerase (XI) expression cassette. This gene, when present in multiple copies, enables alcoholic fermentation of xylose by *S. cerevisiae* (34,35).

## Results and discussion

A consensus delta sequence was obtained by analyzing all SGD entries annotated as delta LTR (**Figure 1a**). A total of 132 sequences between 332 and 338 bp, the most conserved LTRs, were aligned and a consensus 257 bp sequence generated. To compare the selected delta sequence against previously published strategies (**Table S1**), an *in silico* approach was devised. As the main goal of these sequences is to serve as homology templates to guide insertion, the efficiency of a given delta sequence should be proportional to: 1) its length, as longer sequences are more efficient at directing homology directed (36) insertion; and 2) the number of corresponding target sites in the genome, as each additional site increases the odds of an integration occurring. Based on these two criteria, an assortment of delta sequences from the literature was aligned to the yeast reference genome (**Figure 1b**). The delta sequence selected in this work presents the best balance between length of homology and number of right sized hits (257 bp, 85 hits). Shorter homologies, such as those used by Huang *et al*. (32), do not correlate with a greater number of genomic matches (104 bp, 18 hits). Similarly, longer homologies like the ones used Shi *et al*. (31) (408 bp, 8 hits) or the canonical LTR from TyH3 by Boeke *et al*. (18) (334 bp, 17 hits), although longer, also suffer from a relatively small number of hits. Finally, primers annealing the most conserved parts of the alignment were designed and the resulting sequence was amplified and cloned, resulting in plasmid pGU003 (**Figure 1c**). The amplified region was sequenced for confirmation. The δ-integration vector serves as a backbone for multicopy GOI integration, as exemplified next.

**Figure 1.**
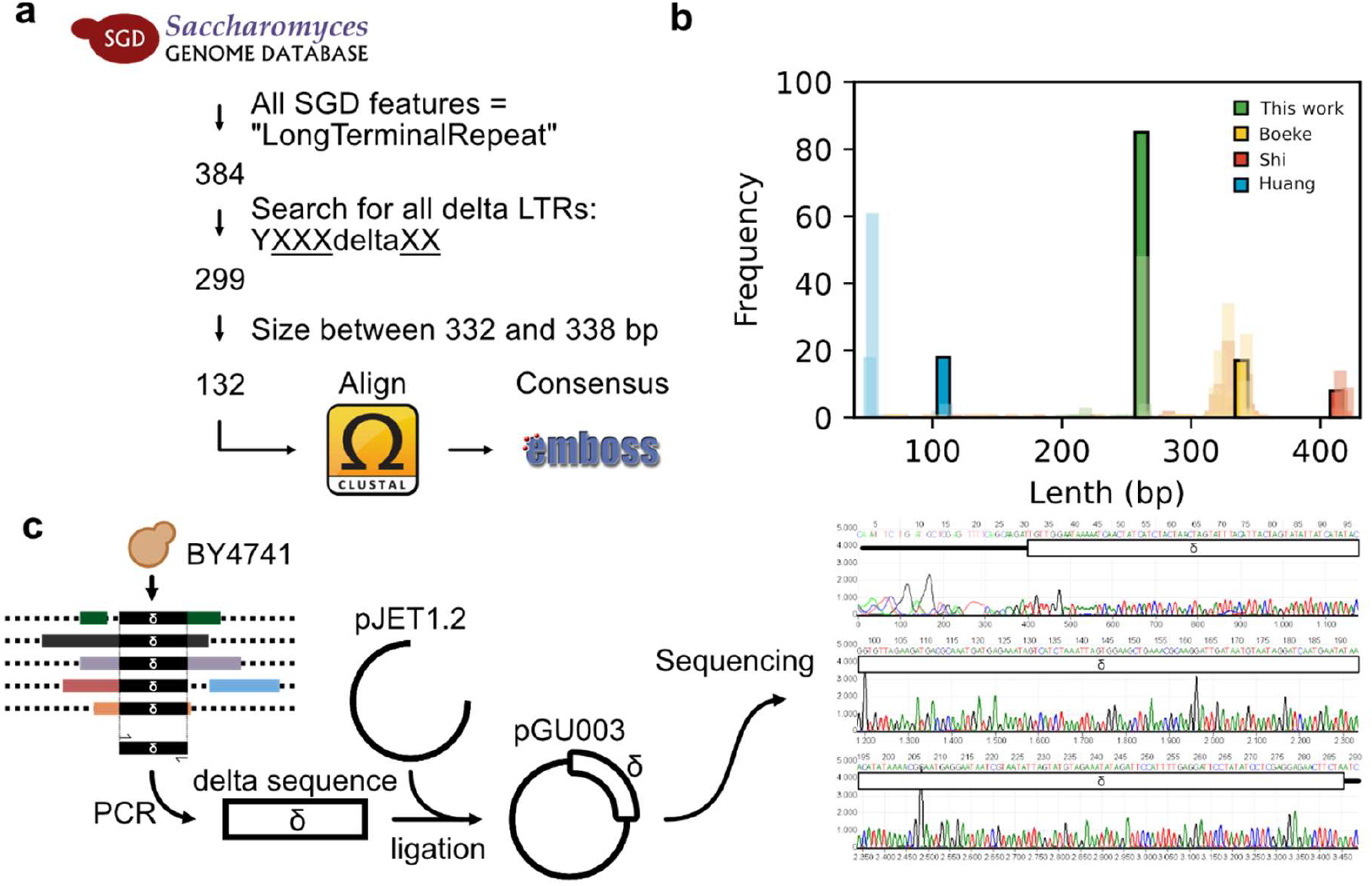
Consensus delta sequence in *S. cerevisiae* for multicopy chromosomal integration. **(a)** Schematic of the *in silico* mining strategy used to identify a conserved delta sequence in *S. cerevisiae*. **(b)** The obtained delta sequence was compared to other sequences from the literature. The results of a BLAST search of the delta sequences against the S288C genome were plotted as a histogram with 1 bp bin size. Each color represents a different delta sequence from literature. Filled bars are the hits with the same length as the query, whereas transparent bars are hits shorter or longer than the intended delta sequence. **(c)** Delta sequence validation and construction of the δ-integration vector pGU003.

To functionally validate the new delta sequence, we sought to introduce the heterologous xylose isomerase (XI) pathway in *S. cerevisiae* to enable pentose fermentation. The xylose isomerase gene is only able to impart robust pentose fermentation when it is present in several copies (35,37–43), therefore a multicopy integration strategy is foremost. Here, the XI expression cassette was amplified from p426_GPD_OrpXI (44) and cloned into pGU003 to obtain pGU003.OrpXI. This plasmid was used as a template to generate donors for genomic integration, containing flanking homology regions of 126/130 bp (left/right) to the delta sequences in the genome. Cells were co-transformed with the donor and a plasmid harboring a KanMX cassette, pGS004, for antibiotic selection of transformants (**Figure 2a**). Delta integration was performed on the yeast BVY270, a robust platform yeast harboring several modifications for pentose fermentation, except a XI (44). The transformed yeasts were plated onto selective media, and antibiotic-resistant colonies were transferred to a 96 well plate. Cells were cultivated 3 times in non-selective media to cure the selection plasmid and then used for a growth assay in xylose. The fermentation kinetics and XI copy number of the best performing colony were further characterized (**Figure 2b**).

**Figure 2.**
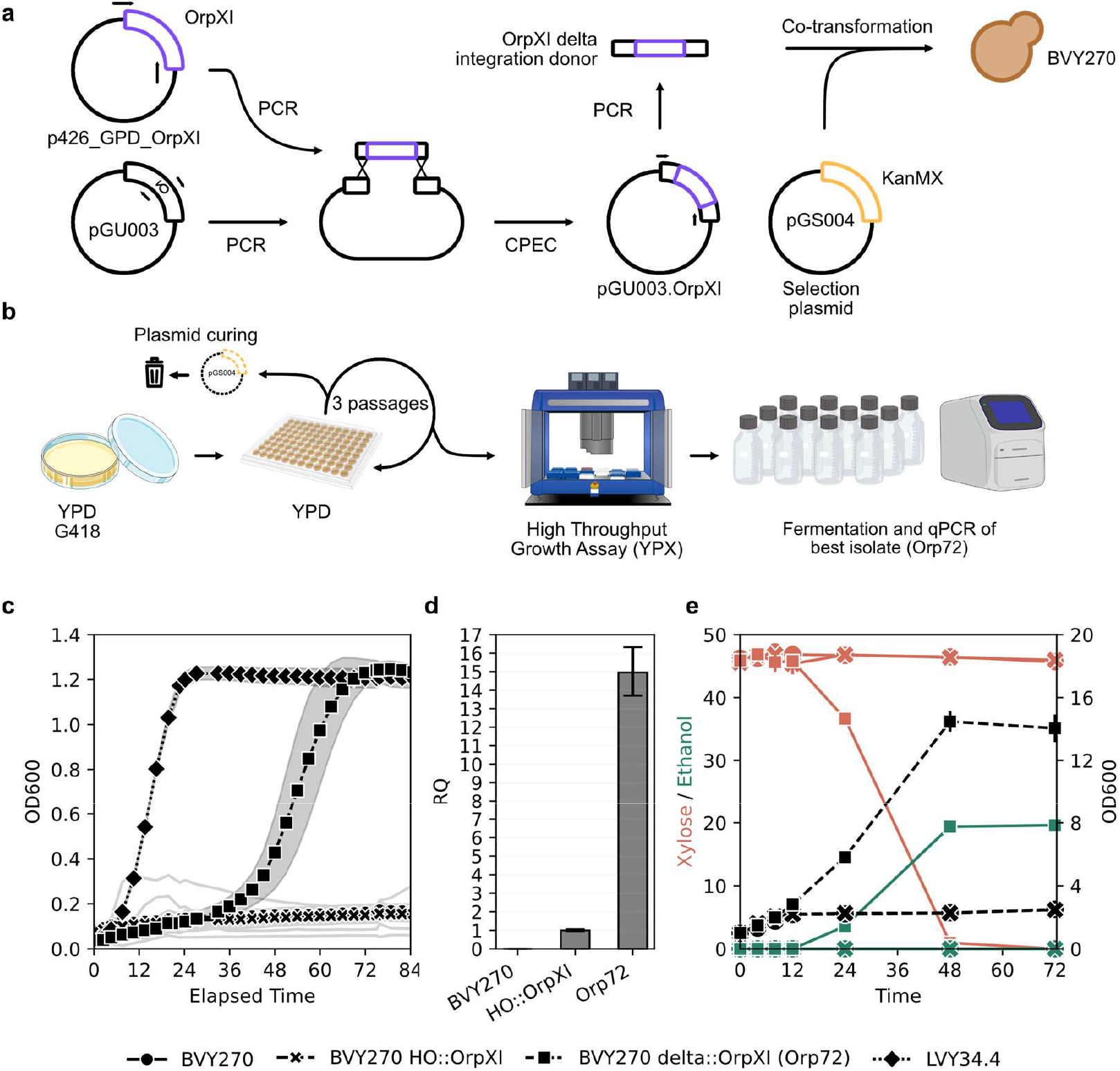
Validation of the novel delta sequence multicopy gene integration strategy by enabling efficient xylose metabolism in *S. cerevisiae*. **(a)** Schematic representation of the XI integration strategy. **(b)** Experimental outline used for characterization of the transformants. **(c)** Growth curves of the 91 transformants in xylose medium at 30ºC. BVY270 is the parental strain, without XI; BVY270 HO::OrpXI expresses only one copy of the XI gene; Orp72 (BVY270 delta::OrpXI) is a transformant from the delta integration and LVY34.4 is a reported strain engineered for xylose metabolism. **(d)** XI copy number determination via qPCR, the parental strain and a control strain with one XI copy integrated into the HO *locus* (HO::OrpXI) were used to determine the XI number of Orp72. **(e)** Fermentation kinetics of the Orp72 strain and BVY270 HO::OrpXI in xylose medium at 30 ºC 250 rpm.

Of the 91 cured colonies assayed for growth in xylose, the colony #72 (Orp72) reached a final optical density (OD600) equal to the positive control, LVY34.4 – an elite industrial strain with 36 integrated copies of XI along with several other mutations obtained after adaptive evolution assays (35). At the same time, strain BVY270 expressing only one copy of the gene (BVY270 HO::OrpXI) could not grow in this medium, revealing once again that multicopy integration is necessary to endow xylose metabolism in *S. cerevisiae* (**Figure 2c**). To quantify the delta integration frequency, 14 cured colonies were randomly picked, and the presence of the XI gene was verified by PCR in 50% of them (7/14) (**Figure S1**). Previous work using the same strategy, but a different delta sequence, achieved an integration frequency of 6.67% (26). Therefore, the observation that only Orp72 was able to robustly grow on xylose, while half the colonies are expected to have an integrated XI, could be explained by a high XI copy number in Orp72. To test this hypothesis, a genomic qPCR for XI was performed on Orp72, revealing 15 integrated copies (**Figure 2d**). Finally, strain Orp72 was evaluated for fermentation performance in medium containing 45.90±0.28 g of xylose (**Figure 2e**). A total of 19,38±0,02 g/L ethanol was obtained in 48 hours, representing a productivity of 0,40±0,001 g/L/h and 0,43±0,003 g/g yield. This represents a remarkable result for xylose fermentation capacity in *S. cerevisiae*, achieved through rational genetic engineering without the use of episomal expression or adaptive evolution assays.

## Conclusion

The rational selection of a conserved delta sequence fragment in *S. cerevisiae* allowed the rapid and facile multicopy integration of a transgene. Up to 15 simultaneous integrations were observed for a single-step transformation. Moreover, this approach proved efficient also for applications requiring less copies, since half of the transformants are expected to harbor at least one copy of the integrative cassette. Transient antibiotic selection using a plasmid borne resistance gene, followed by curing, allows marker-free integration and the reuse of the same antibiotic in downstream engineering steps. The selected delta sequence could be easily amplified from and used in both laboratory (BY4741) and industrial (BVY270 background) yeast strains. We expect that the method provided can accelerate the development of complex pathways involving multiple genes as well as the fine tuning of expression by copy number modulation in yeast.

## Materials and methods

### Consensus delta sequence

All the sequences annotated as “Long terminal repeat” in the *Saccharomyces* genome database (32) were downloaded and filtered for those that contained the keyword “delta” in their systematic name, and which were between 332 and 338 bp in length. They were then aligned using Clustal Omega (45) and a final consensus sequence was build using EMBOSS Cons (46), both on default settings. Primers GUO_056 and GUO_059 were designed to amplify the consensus sequence from BY4741 genomic DNA. This sequence was then cloned into pJET1.2 to obtain pGU003 using the CloneJET PCR Cloning Kit (Thermo Scientific). All primers used in this study are available in **Table S2**.

### *In silico* validation

Previously reported delta sequences were determined whenever possible, either as provided in the manuscript or by using the primers reported to perform *in silico* PCR (47) to recreate the homology sequence described by the authors. The reported delta sequences, as well as the one described here, were searched against the *S. cerevisiae* reference genome using BLAST (48). The resulting hits were plotted and sorted by length. A delta sequence with well-conserved copies in the genome is expected to return homogeneous hits of the same length. Delta sequences less representative should return more diffuse hits of lesser or greater length, indicating deletions and insertions, and therefore poorer homology templates.

### Integrative plasmid

Plasmids for genomic integration into delta sequences were assembled by the circular polymerase extension cloning (CPEC) method (49). Briefly, pGU003 was amplified by inverse PCR using primers that anneal in the middle of the delta sequence (GUO_057 and GUO_058), amplifying the whole vector. The XI expression cassette was amplified from p426_GPD_OrpXI (44) using primers containing homology regions to the delta sequence (BVO_076 and BVO_077). Both amplicons were mixed in equimolar amounts (120 pmol) and assembled *in vitro* by overlap extension using Phusion High-Fidelity DNA polymerase (Thermo Scientific) with the following cycle parameter: 98 ºC 30 s; (98 ºC 10 s; 70 to 55 ºC at 0.5 ºC/s; 55ºC 30 s; 72 ºC 3 min) 25 times; 72 ºC 5 min; 12 ºC hold. *E. coli* DH5α were electroporated with 2 µL of the assembly reaction for storage and replication. All plasmids used in this study are available in **Table S3**.

### Delta integration

The donor sequence, comprised of the expression cassette flanked by homology regions for the delta sequences, was amplified from pGU003.OrpXI with primers GUO_056 and GUO_059. 15 µL of the crude PCR reaction plus 1 µg of pGS004.0 were used to transform yeast cells using the PEG/ssDNA/LiAc/DMSO method described by Gietz et al (50). pGS004.0 is a Cas9 expression plasmid with no sgRNA (51), it was co-transformed alongside the donor to provide a means of selecting competent cells by virtue of its *kanMX* marker. Transformants were collected in YPD (10 g/L yeast extract, 20 g/L peptone and 20 g/L glucose) plates with 200 μg/mL G418 (Gibco). The expression of the Cas9 transgene imposes a metabolic burden, rendering the plasmid unstable, this facilitates the curing process. The rationale, adapted from Guerra *et al*. (26), is that cells which were competent to incorporate the plasmid DNA were also likely to incorporate the linear donor, which was present in great molar excess. For multicopy integration, the strain used was *S. cerevisiae* BVY270 (44). The same strain was transformed with the XI expression cassette targeting the *HO locus* using vector pGS004.29 (51), serving as a control for single copy modified yeast (BVY270 HO::OrpXI). All strains used in this study are described in **Table S3**.

### Growth assay and xylose fermentation

91 transformed colonies of strain BVY270 for delta integration, together with parental BVY270, BVY270 HO::OrpXI (harboring only one copy of XI) and LVY34.4 were cultivated overnight in YPD in 96-well plates at 30 ºC. Cultures were diluted 10 times in YPX (10 g/L yeast extract, 20 g/L peptone and 45 g/L xylose) in a final volume of 150 µL and growth at 30ºC was analyzed using a microplate reader (Spectramax 384 Plus, Molecular Devices). OD600 readings were taken at 180 min intervals for 51 cycles (≈153 h). Next, to validate the fermentation phenotype of Orp72, the strain and control BVY270 HO::OrpXI were cultivated in semi-anaerobiosis in YPX. Fermentation was carried out in 100 mL bottles using 60 mL medium at 30 ºC and 200 rpm, with an initial OD600 of 1. Samples were collected at 0h, 4h, 8h, 12h, 24h, 48h and 72h for high-performance liquid chromatography (Shimadzu LC-2050C 3D) using a Bio-Rad HPX-87H column, 5 mM sulfuric acid as the mobile phase and a flow rate of 0.6 mL/min.

### Quantitative PCR

For qPCR assays, strains were collected after fermentation and genomic DNA was extracted using the Yeast DNA Extraction Kit (Thermo Scientific). Reactions were assembled in 10 µL quadruplicates using the PowerTrack SYBR Green Master Mix (Thermo Scientific) with 10 ng of gDNA. The strain BVY270 HO::OrpXI and actin (ACT1) genes was used as standard. Copy number was determined using the Relative Standard Curve method, a serial dilution of 100, 20, 4, 0.8 and 0.16 ng of BVY270 HO::OrpXI gDNA was used (OrpXI: r^2^=0996 Eff%=100.328; ACT: r^2^=0.997 Eff%=96.74). Primers used are available on **Table S2**.

## Supporting Information

### Supplementary Information

The delta sequences gathered from the literature used for comparison in this study, additional validation of the integration frequency and list of primers, strains and plasmids.

## Acknowledgments

We thank the Laboratory of Genomics and bioEnergy (LGE) and all its members for their advice and support during the investigation. We also acknowledge the São Paulo Research Foundation (FAPESP) for funding this investigation (grant number 2023/02347-0).

## Notes

### Competing Interest Statement

The authors have declared no competing interest.

