## Supplementary Material for "A rational approach for multicopy delta integration in *Saccharomyces cerevisiae* targeting a conserved Ty1 LTR sequence"

Table S1. Delta sequences gathered from the literature and used for comparison in this work. Sequences were used as reported or reconstructed using primers provided by the authors.

| Delta sequence | Origin | Reference |
| --- | --- | --- |
| TGTTGGAATAGAAATCAACTATCATCTACTAACTAGTATTTACATTACTAGTATATTATCATATACGGTGTTAGAAGATGACGCAAATGATGAGAAATAGTCATCTAAATTAGTGGAAGCTGAAACGCAAGGATTGATAATGTAATAGGATCAATGAATATAAACATATAAAATGATGATAATAATATTTATAGAATTGTGTAGAATTGCAGATTCCCTTTTATGGATTCCTAAATCCTTGAGGAGAACTTCTAGTATATTCTGTATACCTAATATTATAGCCTTTATCAACAATGGAATCCCAACAATTATCTCAACATTCACCCATTTCTCA | 3’ LTR from TyH3 cloned by Boeke et al. (1). | Wang, 1996 (2)  Lee, 1997 (143,144)  Guerra, 2006 (3)  Yamada, 2010 (4) |
| TGGAAGCTGAAACGTCTAACGGATCTTGATTTGTGTGGACTTCCTTAGAAGTAACCGAAGCACAGGCGCTACCATGAGAATTGGGTGAATGTTGAGATAATTGTTGGGATTCCATTGTTGATAAAGGCTATAATATTAGGTATACAGAATATACTAGAAGTTCTCCTCGAGGATATAGGAATCCTCAAAATGGAATCTGCAATTCTACACAATTCTATAAATATTATTATCATCATTTTATATGTTTATATTCATTGATCCTATTACATTATCAATCCTTGCGTTTCAGCTTCCACTAATTTAGATGACTATTTCTCATCATTTGCGTCATCTTCTAACACCGTATATGATAATATACTAGTAATGTAAATACTAGTTAGTAGATGATAGTTGATTTCTATTCCAACA | Amplified from strain HZ848 gDNA. | Shi, 2016/2019  (7,8) |
| TCATATACGGTGTTAGACGATGACATAAGATACGAGGAACTGTCATCGGAAGCTGAAACGCAAGGATTGATAATGTAATAGGATCAATGAATATTA | Amplified from strain JUK36α gDNA. Identical to S288c YJRWdelta18 | Huang, 2020 (9) |
| TGTTGGAATAAAAATCAACTATCATCTACTAACTAGTATTTACATTACTAGTATATTATCATATACGGTGTTAGAAGATGACGCAAATGATGAGAAATAGTCATCTAAATTAGTGGAAGCTGAAACGCAAGGATTGATAATGTAATAGGATCAATGAATATAAACATATAAAACGGAATGAGGAATAATCGTAATATTAGTATGTAGAAATATAGATTCCATTTTGAGGATTCCTATATCCTCGAGGAGAACTTCTA | Consensus from all S288c SGD delta sequences. Amplified from strain BY4741 gDNA. | This work |

Table S2. List of primers used in this work.

| Name | Sequence (5’ – 3’) | Use |
| --- | --- | --- |
| GUO_056 | TGTTGGAATAAAAATCAACTATCATCTACTAACTAG | delta1-Fw |
| GUO_057 | GTTTCAGCTTCCACTAATTTAGATGACT | delta1-Rv |
| GUO_058 | GCAAGGATTGATAATGTAATAGGATCAATG | delta2-Fw |
| GUO_059 | CTAGAAGTTCTCCTCGAGGAT | delta2-Rv |
| BVO_076 | ATAGTCATCTAAATTAGTGGAAGCTGAAACTCATTATCAATACTCGCCATT | OrpXI-Fw |
| BVO_077 | CATTGATCCTATTACATTATCAATCCTTGCGCAAATTAAAGCCTTCGAGCGTCCCAAAACCTTCTCA | OrpXI-Rv |
| GUO_092 | AACGGTGCCTCTACTAATCCAG | qPCR-Fw |
| GUO_093 | TAACCTTCTCTACCACCCCAGA | qPCR-Rv |

Table S3. List of strains and plasmids used in this work.

| Strains | Genotype | Reference |
| --- | --- | --- |
| BY4741 | MATa his3Δ1 leu2Δ0 met15Δ0 ura3Δ0 | (11) |
| LVY34.4 | Derived PE-2; MATα; CEN5::pTDH1-xylA-tTDH1; gre3Δ; CEN2::pADH1-XKS1-tADH1; CEN8::pADH1-XKS1-tADH1; CEN12::pTDH1-TAL1-tTDH1-pPGK1-RKI1-tPGK1; CEN13::pTDH1-TKL1-tTDH1-pPGK1-RPE1-tPGK1; pOXylATy1 + adaptive evolution and selection | (12) |
| BVY270 | Derived PE-2; MATα; *CEN5::pTEF1-hphMX6-tTEF1; gre3Δ; CEN2::pADH1-XKS1-tADH1; CEN8::pADH1-XKS1-tADH1; CEN12::pTDH1-TAL1-tTDH1-pPGK1-RKI1-tPGK1; CEN13::pTDH1-TKL1-tTDH1-pPGK1-RPE1-tPGK1* | (13) |
| BVY270 HO::OrpXI | BVY270; *HO::pGPD1-XylA-CYC1t* | This work |
| BVY270 delta::OrpXI | BVY270; *delta::pGPD1-XylA-CYC1t* | This work |
| Plasmids | **Relevant features** |  |
| pJET1.2 | AmpR; ColE1 | Thermo Scientific™ |
| p426_GPD_OrpXI | *URA3*; 2µ; *pGPD1-XylA-CYC1t* | (13) |
| pGS004.0 | pTEF1-Cas9-tCYC1; pSNR52-sgRNA-tSUP4; KanMX | (14) |
| pGS004.29 | pGS004 with gRNA sequence targeting HO in Industrial strains | (14) |
| pGU003 | pJET1.2 delta1-delta2 | This work |
| pGU003.2 | pGU003 delta1-*pGPD1-XylA-CYC1t* -delta2 | This work |

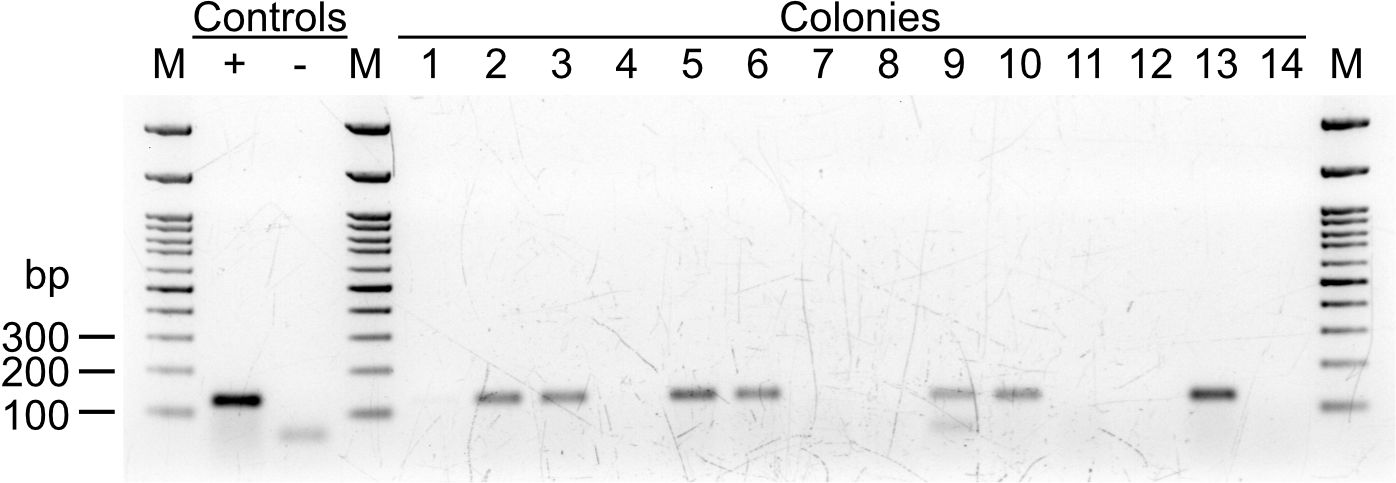

Figure S1. Colony PCR for the OrpXI gene of 14 randomly picked colonies. Half of the colonies assayed displayed the expected 128 bp band. the same primers were used for qPCR. Genomic DNA of BVY270 HO::OrpXI was used as positive control. No template reaction was used as negative control.
